# *tartan* underlies the evolution of male *Drosophila* genital morphology

**DOI:** 10.1101/462259

**Authors:** Joanna F. D. Hagen, Cláudia C. Mendes, Amber Blogg, Alex Payne, Kentaro M. Tanaka, Pedro Gaspar, Javier Figueras Jimenez, Maike Kittelmann, Alistair P. McGregor, Maria Daniela S. Nunes

## Abstract

Male genital structures are among the most rapidly evolving morphological traits and are often the only features that can distinguish closely related species. This process is thought to be driven by sexual selection and may reinforce species separation. However, while the genetic basis of many phenotypic differences have been identified, we still lack knowledge about the genes underlying evolutionary differences in male genital organs and organ size more generally. The claspers (surstyli) are periphallic structures that play an important role in copulation in insects. Here we show that natural variation in clasper size and bristle number between *Drosophila mauritiana* and *D. simulans* is caused by evolutionary changes in *tartan (trn)*, which encodes a transmembrane leucine-rich repeat domain protein that mediates cell-cell interactions and affinity differences. There are no fixed amino acid differences in *trn* between *D. mauritiana* and *D. simulans* but differences in the expression of this gene in developing genitalia suggest cis-regulatory changes in *trn* underlie the evolution of clasper morphology in these species. Finally, analysis of reciprocal hemizyotes that are genetically identical, except for which species the functional allele of *trn* is from, determined that the *trn* allele of *D. mauritiana* specifies larger claspers with more bristles than the allele of *D. simulans*. Therefore we have identified the first gene underlying evolutionary change in the size of a male genital organ, which will help to better understand the rapid diversification of these structures and the regulation and evolution of organ size more broadly.

**Significance Statement:** The morphology of male genital organs evolves rapidly driven by sexual selection. However, little is known about the genes underlying genitalia differences between species. Identifying these genes is key to understanding how sexual selection acts on development to produce rapid phenotypic change. We have found that the gene *tartan* underlies differences between male *Drosophila mauritiana* and *D. simulans* in the size and bristle number of the claspers - genital projections that grasp the female during copulation. Moreover, since *tartan* encodes a protein that is involved in cell affinity, this may represent a new developmental mechanism for morphological change. Therefore, our study provides new insights into genetic and developmental bases for the rapid evolution of male genitalia and organ size more generally.

## Introduction

The morphology of male genitalia can differ dramatically even between very closely related animal species (1). In *Drosophila mauritiana* males, for example, the size, shape and bristle morphology of the claspers (surstyli), posterior lobes (epandrial posterior lobes) and anal plates (cerci) are strikingly different from those of its sister species *D. simulans* and *D. sechellia* (Fig. 1). Moreover, these differences have evolved in only the last 240,000 years since these species last shared a common ancestor (2) (Fig. 1a).

**Fig. 1.**
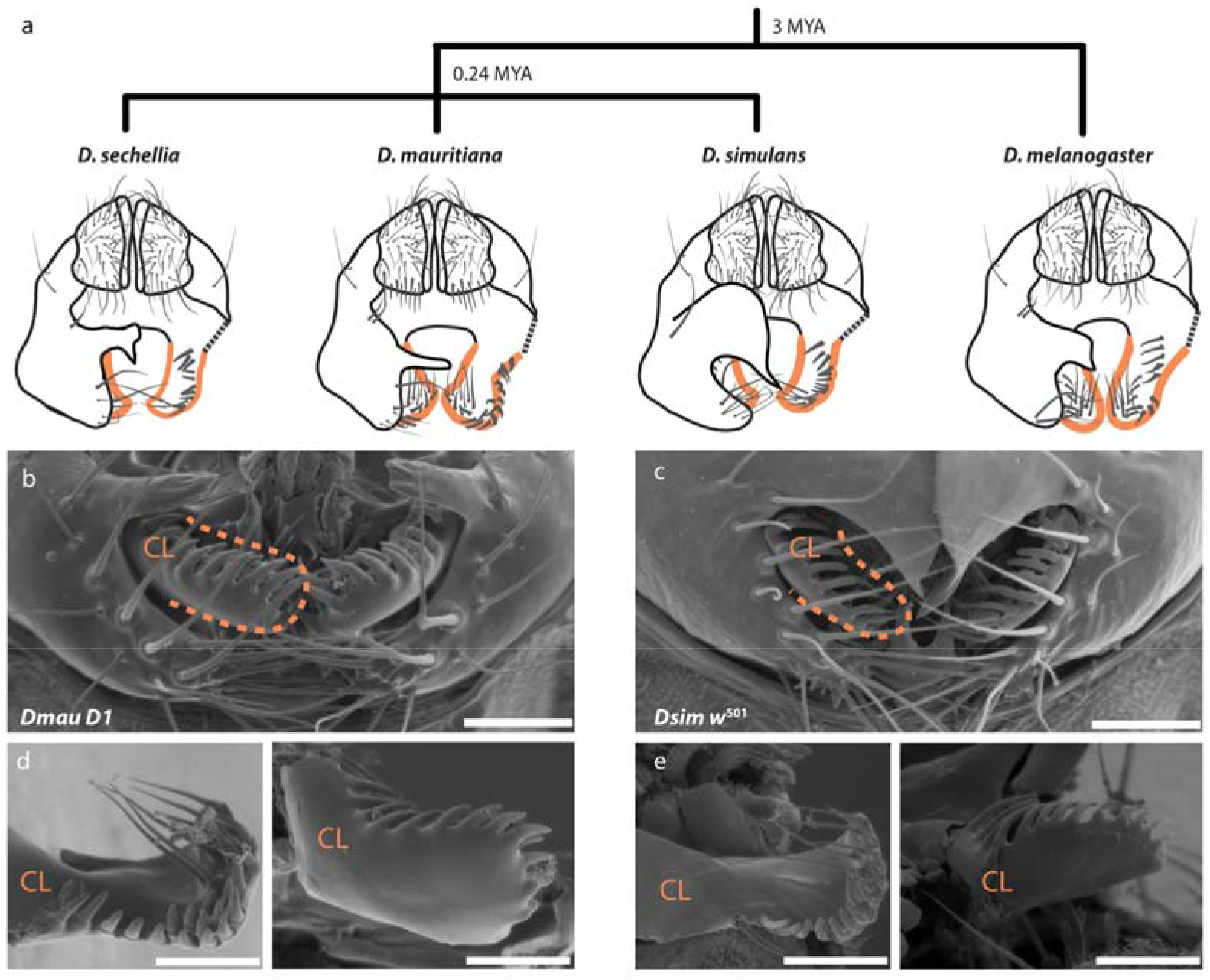
Divergence in periphallic structures in the *D. simulans* clade and its relationship to the outgroup *D. melanogaster* (2). **a.** Schematic representation of the male analia and external genitalia (posterior view). Posterior lobes are illustrated as dissected away on the right-hand-side, in order to facilitate visualisation of the claspers (outlined in orange), which are typically covered by the posterior lobes. While the shape and size of the posterior lobes is species-specific, the claspers and anal plates are very similar between *D. simulans and D. sechellia*, which are smaller and have less bristles than those of *D. mauritiana* and *D. melanogaster*. In addition, the clasper bristles of *D. mauritiana* are shorter and thicker than those of the other three species (19, 20, 57). **b – d**. Scanning electron micrographs of *Dmau D1* (**b & d**) and *Dsimw^501^* (**c & e**) external male genitalia (upper panel) and dissected claspers (lower panels) scale bars = 50 um.

As in other animal groups (1, 3–5), morphological variation of genital structures is thought to have been driven by sexual selection (6), but the mechanism(s) (female choice, sperm competition or sexual antagonism (5)), and its contribution to reproductive isolation between populations and species, has been difficult to address and resolve both theoretically (7–9) as well as experimentally (10, 11). Genetic manipulation of the evolved loci would allow us to test directly the effect of male genital divergence on mating behaviour and reproductive fitness and therefore facilitate the empirical study of these questions (12, 13). However, although quantitative mapping studies of morphological differences in male genitalia between species of the *D. simulans* clade were first carried out more than three decades ago (14–21), the genetic basis of male genital divergence between these species has remained elusive. This is due, at least in part, to the large number of loci found to contribute to variation in size and shape of these structures (18, 19, 21).

The claspers are periphallic structures with an essential role in grasping and proprioception of the female, and in securing genital coupling (12, 22–27). Previously, we found that multiple loci contribute to variation in clasper size and bristle number between *D. simulans* and *D. mauritiana* (19). Here we describe mapping and functional experiments that strongly suggest that cis-regulatory changes in *trn* underlie differences in clasper morphology between these two species.

## Results and Discussion

Previously, we identified two regions on the left arm of chromosome 3 that contribute to differences in clasper size and bristle number between *D. mauritiana* and *D. simulans* (19). Here, we have generated new recombinant introgression lines (ILs) between the *D. mauritiana D1 (Dmau D1)* and *D. simulans w*^501^ *(Dsim w*^501^) strains (Supplementary File 1) to increase the resolution of one of these regions, C2, from approximately 3.5 Mb (24) to 177 kb. This interval explains about 16.3% of the difference in clasper size (and 37.9% of clasper bristle number) between the two parental strains (Fig. 2 and Supplementary File 2a and 2b). The claspers of lines that are homozygous for introgressed *D. mauritiana* DNA in C2 are larger than those of natural strains of *D. simulans* (Extended Data Fig. 1). The change in clasper size caused by differences in C2 is therefore outside the range of variation in clasper size in *D. simulans*, suggesting that C2 underlies interspecific divergence between *D. mauritiana* and *D. simulans* and not merely intraspecific polymorphism in clasper size in either or both of these species.

**Fig. 2.**
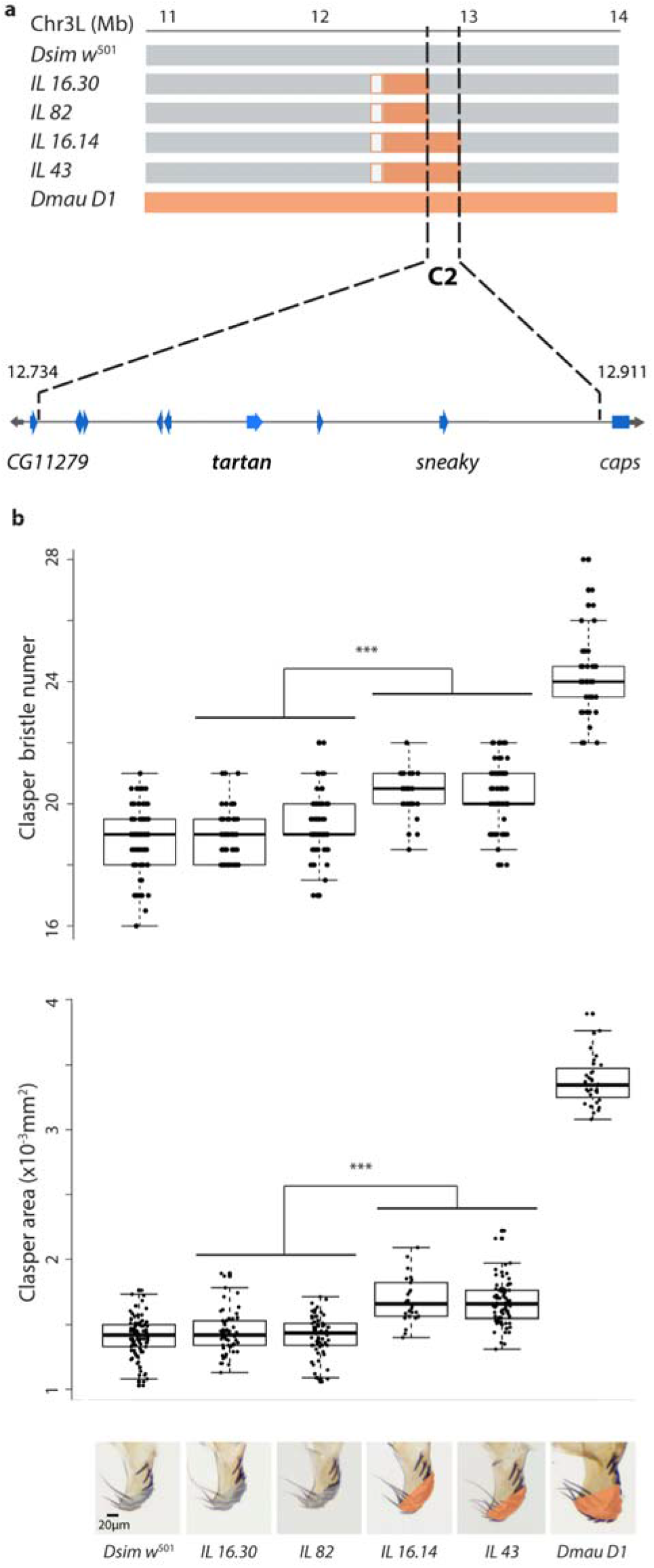
High-resolution mapping of differences in clasper morphology between *Dsim w*^501^ and *Dmau D1*. **a.** Introgression line breakpoints on chromosome arm 3L define the 177 kb region C2 (grey, orange and white boxes indicate DNA regions from *Dsim w*^501^, *Dmau D1* or not verified, respectively). Coordinates are given in Mb with respect to the *D. simulans* genome (Flybase R2.02). This region contains eight protein coding genes including *trn* and is flanked by *CG11279* and *caps*. **b.** Introgression lines containing region C2 from *Dmau D1 (IL43* and *IL16.14*) contribute 37.9% of the difference in bristles (upper graph) and 16.3% of the clasper size (lower graph) difference of this strain compared to *Dsim w*^501^ (Supplementary File 2a). *IL43* and *IL16.14* differed significantly from *IL16.30* and *IL82* in clasper bristle number and in clasper area (*p* < 0.001). Asterisks indicate significance comparisons where *p* < 0.001 (Supplementary File 2c). Shading in the bottom panel indicates the area measured at the distal end of the claspers in lines containing *Dsim w*^501^ (grey) or *Dmau D1* regions (orange) for C2. Boxes indicate the range, upper and lower quartiles and median for each sample.

C2 contains eight protein-coding genes with orthologs in *D. melanogaster*. RNA-Seq data suggests that only one of these genes, *tartan (trn)*, is expressed in the terminalia of *D. simulans* and *D. mauritiana* when the difference in clasper morphology develops between these two species (Supplementary File 3 and Extended Data Figs. 3). However, if the causative gene has a very localised pattern of expression its expression may not have been detected in the RNA-Seq. Therefore, we knocked-down the expression of all genes in the candidate region (with the exception of *CG34429*, for which there was no available UAS line) using RNAi in *D. melanogaster* to test if these positional candidates are involved in clasper development (Supplementary File 4). In addition, we knocked-down *CG11279* and *capricious (caps)* – a gene that also encodes a leucine-rich repeat transmembrane protein closely related to *trn* and that functionally overlaps with *trn* in some contexts (28–34). These two genes flank C2, but their cis-regulatory sequences may still be within this region (Fig. 2a). We found that while knockdown of *trn* significantly reduced the size of the claspers (Supplementary File 4 and Extended Data Fig. 2), RNAi against any of the other nine genes tested, including *caps*, had no effect on clasper morphology in *D. melanogaster* (Supplementary File 4). Note that *trn* RNAi had no effect on the posterior lobes consistent with region C2 only affecting the claspers (Supplementary File 4).

It is thought that the main function of *trn* is to confer differences in affinity between cells and mediate their correct allocation to compartments in developing tissues such as the nervous system, trachea, eyes, wings and legs (28, 30, 32, 35–37). Intriguingly, changes in *trn* expression can affect the allocation of cells between compartments, cause misspecification of compartmental boundaries, and even result in invasive movements of cells across such boundaries (33, 36, 37). Our RNA-Seq data indicates that *trn* is more highly expressed in *D. simulans* during early terminalia development, but is subsequently up-regulated in *D. mauritiana* at a later stage (Supplementary File 3). However, these data correspond to the sum of all the expression domains of *trn* throughout the terminalia at each of these stages and may conceal more subtle localised expression differences between these species in specific tissues like the developing claspers. Therefore, we investigated the spatial pattern of *trn* expression throughout terminalia development using mRNA *in situ* hybridisation (ISH) in *Dmau D1* and *Dsim w*^501^ (Fig. 3a-c and Extended Data Fig 3). Concomitantly, we observed a four hour difference in the timing of terminalia development between the two strains used (Fig. 3a-c; Extended Data Fig. 3). We observed that during early pupal stages *trn* is more highly expressed in *Dsim w*^501^ compared to *Dmau D1* at the centre of the terminalia, from where the internal genital structures will develop, which may explain the overall higher expression of *trn* in *D. simulans* at 30 hAPF according to the RNA-Seq data (Fig. 3a and 3b; Supplementary File 3). However, during later stages, the expression of *trn* is detected in a wider domain and persists for longer at the base of the developing claspers of *Dmau D1* compared to *Dsim w*^501^ (black arrowheads in Fig. 3a and 3b), consistent with higher expression of *trn* in *D. mauritiana* detected in the RNA-Seq data at approximately 50 hAPF (Supplementary File 3). These results are also consistent with the RNAi results in *D. melanogaster* where knockdown of *trn* results in the loss of *trn* expression at the base of the claspers (Extended Data Fig. 2b) and the development of smaller claspers (Extended Data Fig. 2a). Together, these results suggest that the higher and/or more persistent expression of the *trn^mau^* allele relative to the *trn^sim^* allele in the developing claspers is responsible for the larger claspers in *D. mauritiana*.

**Fig. 3.**
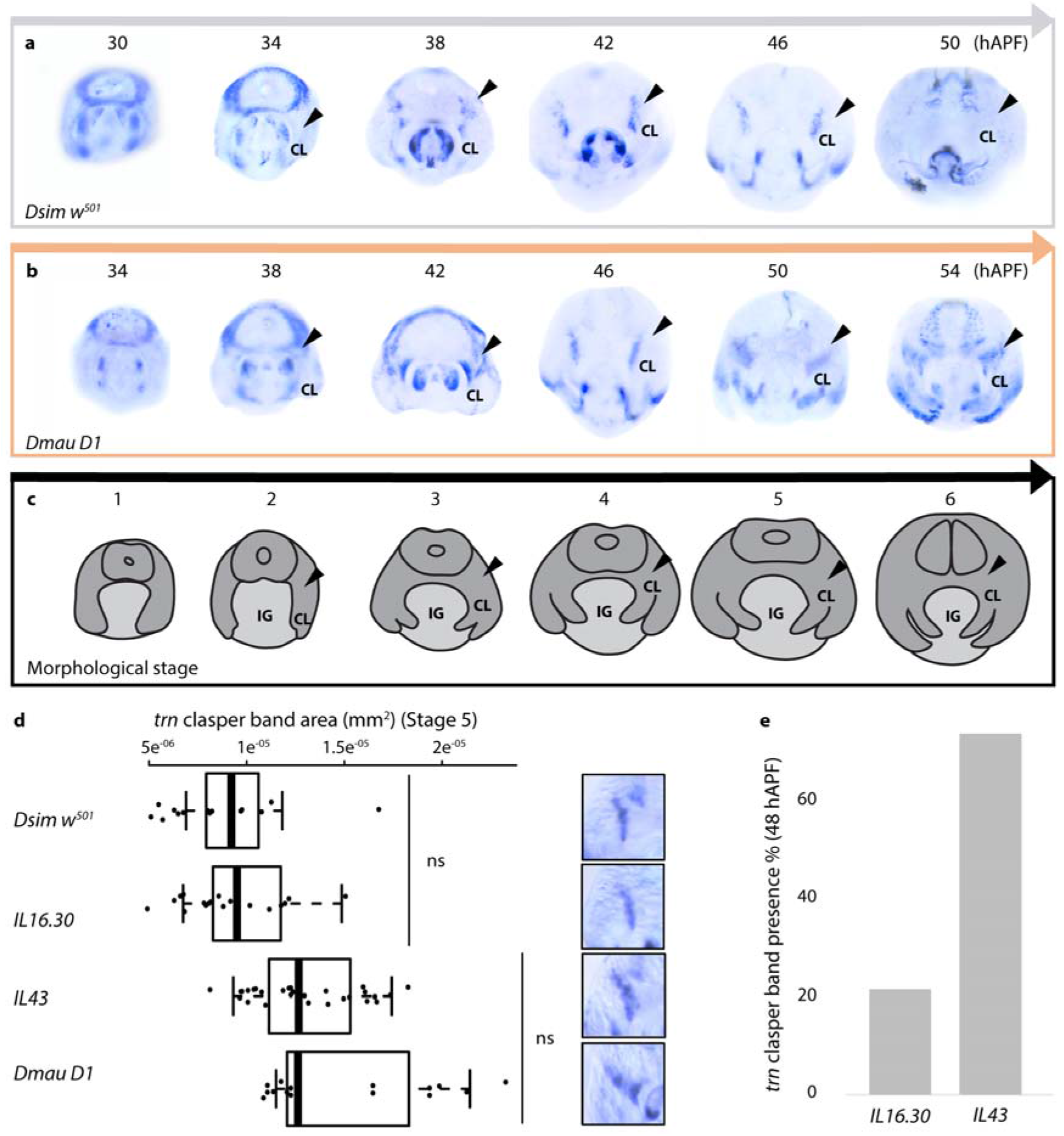
The spatial and temporal expression of *trn* differs in the developing claspers of *D. simulans* and *D. mauritiana*. Expression shown at four hour intervals hours after puparium formation (hAPF) in *Dsim w*^501^ (**a**) and *Dmau D1* (**b**). **c**. Illustration of the developing structures at each morphological stage (Extended Data Fig. 3). Black arrowheads indicate expression at the base of the developing claspers. IG: Internal Genitalia; CL: Clasper. (**d**) Analysis of *trn* expression domain at the base of the developing clasper at Stage 5. *trn*^sim^ males, *Dsim w*^501^ and *IL 16.30*, exhibit significantly smaller expression domains than *trn*^mau^ males, *IL 43* and *Dmau D1* (all comparisons in *trn* expression domain between lines are significant (*p* < 0.001), except for those indicated by the brackets, see also Supplementary File 5b). Boxes show the range, upper and lower quartiles, and the median for each sample. Representative *trn* expression at the base of the claspers is shown on the right-hand side of the panel. (e) The proportion of males with *trn* expression at the base of the clasper at 48 hAPF (between stages 5 and 6) in *IL 16.30* and *IL 43*. 51.9% more *IL 43* males exhibit *trn* expression at the base of the claspers compared to *IL 16.30* males (Supplementary File 5c).

Quantitative analysis of *trn* ISH confirmed that males containing *trn^mau^: Dmau D1* and *IL43*, exhibit a larger expression domain at the base of the developing claspers at stage 5 (50 hAPF for *Dmau D1* and 46 hAPF for *Dsim w*^501^ and ILs) than those containing *trn^sım^: Dsim w*^501^ and *IL16.30* (Fig. 3d and Supplementary File 5a). Moreover, although at stage 6, *IL43* and *IL16.30* seem to recapitulate the pattern observed in *Dsim w*^501^ (i.e. *trn* expression no longer detected, *data not shown)*, we found that just before this, between stages 5 and 6 (48 hAPF in these ILs and *D. simulans*, Extended Data Fig 3), there was variability in the presence of *trn* expression at the base of the developing claspers: expression was observed in 21% of *IL16.30* males (i.e. males with *trn^sım^)*, while in 74% of *IL43* males (i.e. males with *trn^mau^)* (Fig. 3e and Supplementary File 5b). These data further supports the hypothesis that spatial and/or temporal divergence in the expression of *trn* underlies differences in clasper size between *D. simulans* and *D. mauritiana*.

We also carried out ISH for *CG11279* and *caps* (which are both also expressed in the terminalia, Supplementary File 3) and *CG34429* (which we were unable to knockdown in *D. melanogaster*). This showed that, unlike *trn*, these genes are either not expressed in the developing genitalia or not in a pattern consistent with a role in clasper development and evolution (Extended Data Fig. 4). For example, although *caps* expression in the male genitalia is generally similar to that of *trn, caps* transcripts were never detected at the base of the developing claspers (Fig. 3a and 3b and Extended Data Fig. 4a).

There are a total of 22 nucleotide differences in the coding sequence of *trn* between our mapped strains, *Dmau D1* and *Dsim w*^501^, and only three of these are non-synonymous (Supplementary File 6). Although none of these substitutions are fixed between the two species, they could be responsible for the difference in clasper size between the two strains used in this study. However, comparison of clasper size between strains of *D. simulans* and *D. mauritiana* with different combinations of amino acids at these three sites suggests that none of them contributes to the difference in clasper size between *D. mauritiana* and *D. simulans* (Extended Data Fig. 5 and Supplementary File 2e). Furthermore, the clasper size of the two mapped strains is well within the range of their corresponding species (Extended Data Fig. 1 and Supplementary File 2d). Taken together our mapping, RNAi in *D. melanogaster*, and expression analysis in developing claspers, suggest that it is most likely that cis-regulatory changes in *trn* underlie differences in clasper morphology between *D. mauritiana* and *D. simulans*.

To confirm that sequence divergence in *trn* contributes to the difference in clasper morphology between *Dmau D1* and *Dsim w*^501^, we used CRISPR/Cas9 to make null alleles of *D. simulans trn* (in *Dsim w*^501^) and *D. mauritiana trn* (in *IL43*, see Fig. 2; Extended Data Fig. 6). We then generated reciprocal hemizyotes for *trn* i.e. genetically identical male flies that differ only in whether they have a functional copy of *trn* from *D. mauritiana* or *D. simulans* (Fig. 4a) (38). Comparison of the claspers between male reciprocal hemizyotes of *trn* shows that flies with a functional *D. mauritiana trn* allele have significantly larger claspers *(p* < 0. 001) with more bristles (*p* < 0.05) than those with a functional *D. simulans trn* allele (Fig. 4 b, Supplementary File 2c). This confirms that, consistent with the effects of the introgressions containing *trn* (Fig. 2), the *D. mauritiana trn* has evolved to confer larger claspers than *D. simulans trn*.

**Fig. 4.**
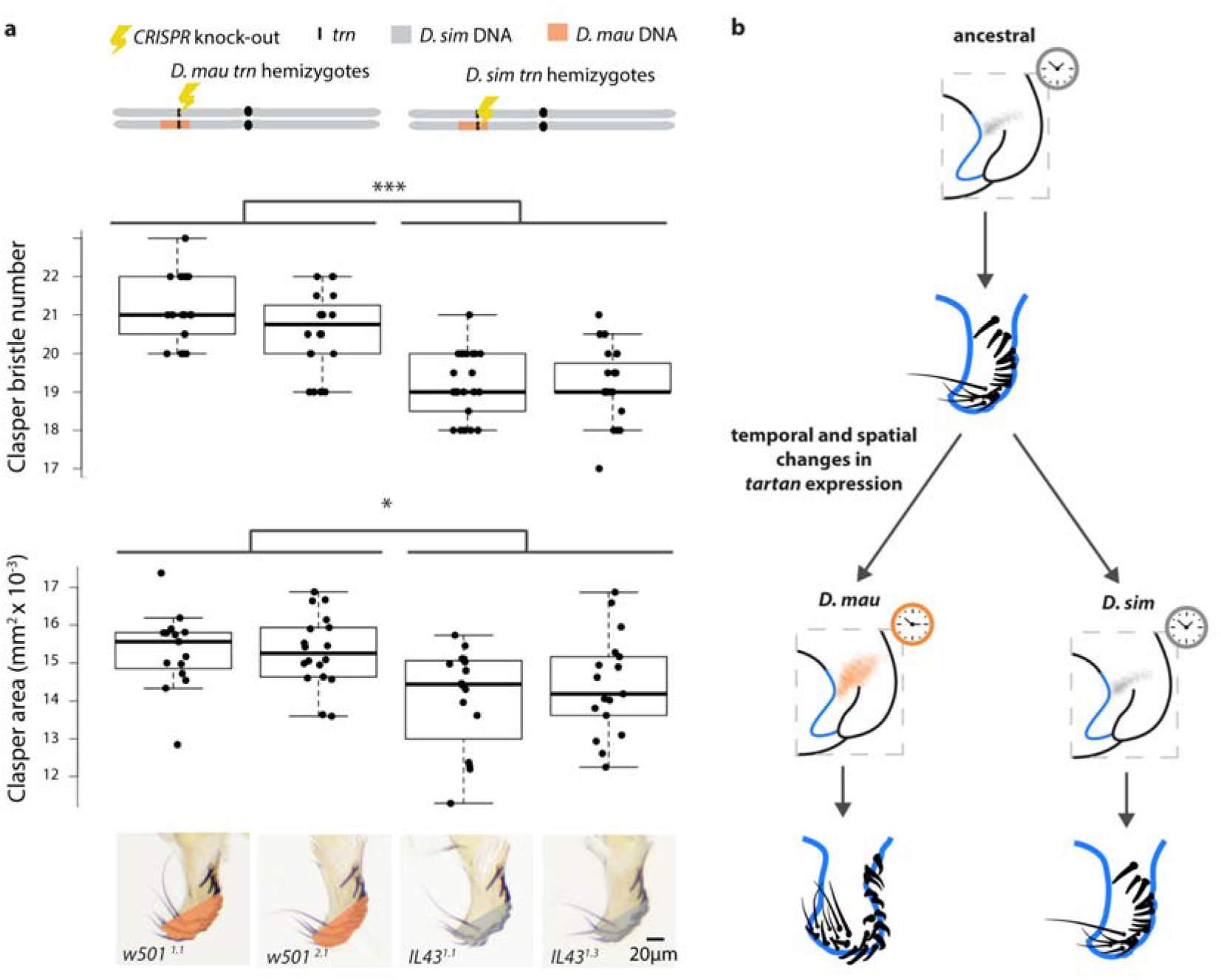
Reciprocal hemizygotes of *trn* show that this locus contributes to evolutionary differences in male clasper morphology. **(a)** Schematic at the top illustrates the 3^rd^ chromosome of the reciprocal hemizygotes carrying a functional allele of *trn* from only *Dmau D1* (left) and *Dsim w*^501^. We found a significant difference in their clasper area (F_(3, 61)_ = 7.012, *p* < 0.001) and clasper bristle number (F_(3, 83)_ = 26.29, *p* < 0.001), shown in the boxplots underneath. Flies with a functional *trn* allele from *D. mauritiana* (*IL43^1.1^* and *IL43^1.3^*), have significantly larger claspers (\*\*\**p* < 0.001) with more bristles (\**p* < 0.05) than those with a functional *D. simulans trn* allele, *w501*^1.1^ and *w501*^2.1^ (Supplementary File 2d). Boxes show the range, upper and lower quartiles, and the median for each sample. **(b)** Evolutionary changes increased the spatial domain and temporal expression of *trn* during clasper development in *D. mauritiana* have led to larger claspers with more bristles in this species compared to *D. simulans*. Orange and grey shading indicate broad and narrow expression of *trn* at the base of the developing claspers in *D. mauritiana* and *D. simulans* respectively. The correspondingly coloured clocks indicate differences in the persistence of this expression domain.

*trn* is the first gene to be identified that underlies the rapid evolution in the size of a male genital organ and more generally one of the first loci found to contribute to natural variation in animal organ size (e.g. 39, 40, 41). While there are many examples of phenotypic evolution caused by changes in the expression of transcription factors and signalling molecules (42), including differences in genital bristles between other *Drosophila* species (43), *trn* encodes a leucine-rich repeat domain transmembrane (28, 30, 32, 33, 36, 44). *trn* appears to mediate affinity differences in cell-cell contact directly through its extracellular domain, directing mispositioned cells towards cues that are currently unknown (33, 36). Our results suggest that differences in *trn* expression in *Drosophila* are able to alter clasper size. Therefore, changes in cell affinity caused by variation in the temporal and/or spatial expression of transmembrane proteins that mediate cell affinity may represent a new mechanism for the evolution of organ size. However, there is also some evidence that *trn* could act as a ligand and may transduce signals, although its intracellular domain appears to be dispensable for most of its functions (28, 30, 33, 44). Therefore, further study into the function of *trn* and characterisation of its role in organ size regulation and evolution is required.

## Materials and Methods

### Introgression mapping and phenotyping

We generated new recombinants from introgression line *D11.01*, which contains *D. mauritiana w^-^ (Dmau w^-^)* DNA in the genomic location 3L:7527144… 15084689 Mb, encompassing the candidate regions C1 and C2 (24). To increase the resolution of the candidate region C2 (19) we backcrossed virgin *D11.01/Dsim w*^501^ heterozygous females to *Dsim w*^501^ males and selected against the visible marker D1 (19, 45) but retained *D. mauritiana* DNA in the predicted C2 region by genotyping with molecular markers (Supplementary Files 1 and 7). Novel recombinants were identified using restriction fragment length polymorphisms (RFLPs) and then maintained as homozygous stocks. Flies were phenotyped and genotyped as described previously (19), using molecular markers (Supplementary File 7). All stocks and crosses were maintained on a standard cornmeal diet at 25°C under a 12-h:12-h dark/light cycle unless otherwise stated.

The posterior lobes were dissected away from the claspers and anal plates, and T1 legs were also retained. The claspers and T1 tibia were mounted in Hoyer’s medium, and images were taken using a Zeiss Axioplan light microscope at X250 magnification for the claspers and X125 for the T1 legs, using a DFC300 camera. Clasper area (see shaded area in Fig. 2b) and tibia length were measured manually using ImageJ (46), and bristle number counted for each clasper. T1 tibia length was used as a proxy for body size, in order to control for the consistency in rearing conditions. Most introgression lines showed no difference in T1 tibia length (Supplementary File 2a), and since genitalia are hypoallometric (13, 17, 18, 47–49), the phenotypic data was not further corrected for body size.

We first tested the normality of the introgression lines duplicates. Depending on the result of this analysis, we conducted either a Kruskal-Wallis followed by a Wilcoxon rank sum test, or an ANOVA followed by a Tukey’s test in order to determine any significant differences between duplicates. If duplicates were not significantly different from each other, the phenotypic measurements were combined. We then compared the phenotype of each the introgression lines to the parental *Dsim w*^501^ strain using a Dunnett’s test (Supplementary File 2a). Region C2 was determined by conducting a Kruskal-Wallis (X^2^ = 92.4, *p* < 0.001, df = 3) followed by a Wilcoxon rank sum test (clasper bristle number) and an ANOVA (F(5, 219) = 42.9, *p* < 0.001) followed by Tukey’s test (clasper area) between *IL 43, IL16.14* and *IL16.30, IL82* (Supplementary File 2b). The effect of introgression lines was calculated as a percentage of the difference between the parental *Dsim w*^501^ and *Dmau D1* strains and was averaged over all lines used to map C2 to determine final effect size (Supplementary File 2a). All statistical analyses were conducted in R Studio. Raw phenotypic data is available in Supplementary File 2f.

### Scanning electron microscopy

For intact male *Drosophila* genitalia, the fly heads were removed and flies placed into fixative (2% PFA, 2.5% GA in 0.1 M NaCac buffer) for 2 hours. To visualise the claspers, genitalia were dissected in Hoyer’s and then placed into fixative. Samples were washed in water and fixed in 1% Osmium over night at 4°C. Osmium was removed, flies washed with water and then taken through a series of ethanol dilutions up to 100% ethanol for dehydration. After 24 hours of 100% ethanol, flies were processed in a critical point dryer and mounted to SEM stubs with Conductive Silver Epoxy (Chemtronics) or coated carbon tabs and gold coated for 30 seconds using a sputter coater. Genitalia were imaged at 2560 × 1920 px in a Hitachi S-3400N SEM in SE mode at 5 kV. Working distance ranged between 7 and 10 mm.

### RNA sequencing and differential expression analysis

We generated three independent biological replicates of RNA-Seq libraries for *Dsim w*^501^ and *Dmau w^-^* terminalia. Males were collected at the white pupal stage by sorting gonad size and placed in a humid chamber, and dissected at 30 (from before any obvious indication of clasper development) and 50 hAPF (near the end of clasper development) (Tanaka et al., unpublished; 50)). Because early pupal tissues are soft, we flash froze the whole pupae by placing on cooled aluminium block with a cake of dry ice. Abdominal tips from 20–30 males were collected to extract the total RNA per biological replicate. The total RNA was extracted using TRIzol Plus RNA Purifica-tion Kit (Life Technologies). The samples were DNaseI (Invitrogen) treated to avoid DNA contamination and the RNA quality was checked using TapeStation (Agilent Technologies). Using 300 ng of total RNA, indexed libraries were generated using the combination of KAPA Stranded mRNA-Seq Kit (KAPA Biosystems) and Adapter Kit (FastGene). Indexed libraries were sent to the Macrogen Japan for sequencing in single lane of HiSeq4000 (Illumina), producing 100 bp paired-end reads. Raw fastq files were quality controlled by FastQCs (ver. 1.34) with the following criterion: minimum length 50 bp, the average Q-score > 20, and continuous base “N” < 2. Filtered reads were mapped to reference coding sequence (CDS) set from (51) using Bowtie2 (ver. 2.2.9). Read counts per CDS were extracted using Samtools (ver. 1.3.1) and the reads per kilo base per million mapped reads (RPKM) were calculated. Raw fastq files are deposited at DDBJ under the accession numbers DRA006755 and DRA006758 for *D. mauritiana* and *D. simulans*, respectively. Genes were considered not to be expressed if RKPM was below 1.5. RNA-Seq analysis of genes in C2, *CG11279* and *caps* is summarised in Supplementary File 3.

### RNAi knockdown of C2 candidate genes

We conducted an RNAi knock down of all the genes within region C2 (with the exception *CG34429* for which there was no available UAS line) in *D. melanogaster* using UAS-RNAi lines from both Vienna Drosophila RNAi Center and TRiP lines from Bloomington stock center (Supplementary File 8). For raw phenotypic data, see Supplementary File 2a. UAS males of our candidate genes were crossed to female NP6333–GAL4 driver virgins (P(GawB)PenNP6333) (52) carrying the transgene UAS-Dicer-2 P(UAS-Dcr-2.D). Crosses for the RNAi were carried out at 25°C. The genital morphology of the male knockdowns was compared to NP6333-Gal4; UAS-Dicer and UAS-RNAi controls. Clasper bristle number and tibia length were measured for 16 individuals of each genotype. Differences in clasper bristle number and tibia size were assessed using a one-way ANOVA followed by a Tukey’s test (Supplementary File 4).

### *trn* sequence analysis

To evaluate if any of the nucleotide differences in the coding sequence of *trn* were fixed between species we took advantage of two population datasets available for *D. simulans* and *D. mauritiana*. One of these datasets consists of Pool-seq data from 107 strains of *D. mauritiana* and from 50 strains of sub-Saharan *D. simulans* (53, 54) available at http://www.popoolation.at/pgt/. To compare allele frequency at the same sites between the two Pool-seq datasets we used a script, kindly provided by Ram Pandey, that aligns the genomes of both species using MAUVE (55) and retrieves the corresponding coordinates and allele frequency information. The data for the coding sequence of *trn* is shown in Supplementary File 6. The other dataset consists of whole genome data for ten strains of each species submitted to the SRA database by the University of Rochester *(D. mauritiana* lines: SRX135546, SRX688576, SRX688581, SRX688583, SRX688588, SRX688609, SRX688610, SRX688612, SRX688710, SRX688712; *D. simulans* lines: SRX497551, SRX497574, SRX497553, SRX497563, SRX497558, SRX497564, SRX497559, SRX495510, SRX495507, SRX497557). An alignment of the *trn* region was kindly provided by the Presgraves lab and is included in the Supplementary File 9. This file also includes the sequences of *D. simulans Kib 32* and *D. mauritiana MS17* (extracted from the Pool-seq data mentioned above) as well as *Dsim w*^501^ and *Dmau D1*, which were resequenced using the primers trn1 – trn4 listed in Supplementary File 7. The sequence analysis is summarized in Supplementary File 6.

To assess the association between each of three non-synonymous amino acid differences and clasper divergence between our mapped strains, we measured clasper size and clasper bristle number of *D. simulans* and *D. mauritiana* strains with different combinations of the three non-synonymous amino acid differences using the methodology described above. For raw phenotypic data, see Supplementary File 2f.

### In situ hybridisation

Staged male pupae which had been incubated at 25°C were flash-frozen on a metal heat block cooled to −80°C. The posterior third of the pupae were cut off and fixed in 4% paraformaldehyde/PBT for half an hour, and washed in methanol and stored at −20’C. Before the *in situ*, the vitelline membranes were peeled away in ice cold methanol. Note that the stages collected in *D. simulans* and *D. mauritiana* are different, due to our observation that our *Dmau D1* strain develops approximately 4 hours more slowly than *Dsim w*^501^ (Extended Data Fig. 3).

Total RNA was extracted from *Dmau D1, Dsim w*^501^ and *D. melanogaster* w^1118^ at a range of developmental time-points using Trizol extraction. A Quantitect Reverse Transcription Kit (Qiagen) was used to synthesize cDNA, in which gene-specific fragments were amplified separately for each species. Primers for *trn, CG11279, CGCG34429* and *caps* were designed using Primer3 (http://primer3.ut.ee) with the addition of T7 linker sequences added to the 5’ end of each primer. To FWD primers (sense) and REV primers (antisense) we added ggccgcgg and cccggggc respectively. Primers are as follows; *trn* (514 bp) ATCGAGGAGCTGAATCTGGG and TCCAGGTTACCATTGTCGCT, *CG11279* (458 bp) CATCTCGAAGTCGGTCAACA and AGGGTCACCTGACCATCAAT, *CGCG34429* (393 bp) GGCTTTGGTATACCTGCAGAA and TGAGCAGGATGTGAAGCACT and *caps* (520 bp) CCGGGAGAACTAACCTTCCA and CTTCATCCAGGCTGCTCAAC. Probes were labelled with 10x DIG labelling mix (Roche Diagnostics) and T7 RNA polymerase (Roche Diagnostics). Purple/dark blue staining was detected using alkaline phosphatase-conjugated anti-DIG antibody FAB fragments (Roche Diagnostics) and Nitro Blue tetrazolium/5-bromo-4-chloro-3-indolyl-phosphate NBT/BCIP (Roche Diagnostics). In situ hybridizations were based on the Carroll lab “Drosophila abdominal *in situ*” protocol (http://carroll.molbio.wisc.edu/methods.html) with minor modifications.

### Quantifying temporal and spatial *trn* expression

To investigate potential differences in *trn* expression domain between introgression lines used to map the C2 interval, in situ hybridisations were carried out using the above methodology at 46 hAPF in *Dsim w*^501^, *IL 16.30, IL 43*, and 50 hAPF in *Dmau D1* (this is the time point that the most profound differences in *trn* expression could be visually detected between the parental species (see Fig. 3a and 3b), and lines were determined to be morphologically equivalent at these stages, see Extended Data Fig. 3). Samples were grouped by the timing the stain was left to develop, mounted in 80% glycerol and imaged using a Zeiss Axioplan light microscope at X125 magnification. Genital arch area and the area of *trn* clasper expression domain were manually measured blind using ImageJ (46). For raw measurements of the *trn* clasper expression domain, see Supplementary File 5c. Differences in the size of the *trn* expression domain was assessed using a one-way ANOVA followed by a Tukey’s test (Supplementary File 5a).

Temporal differences in *trn* expression were tested by recording the presence/absence of expression at the base of the clasper following *trn* in situ hybridisation (see above in situ hybridisation protocol) in *IL 16.30* and *IL 43* at 48 hAPF. Samples were mounted in 80% glycerol and the presence or absence of *trn* at the base of the clasper was manually counted for 25 – 30 samples per line (Supplementary File 5b).

### Generation of reciprocal hemizygotes and statistical analysis

We generated a double-stranded cut 121 bp into the first coding exon of *trn*, resulting in a frameshift mutation. The cut was mediated by pCFD3 gRNA plasmid and pHD-DsRed-attp donor plasmid co-injected by The University of Cambridge Department of Genetics Fly Facility at 0.1ug/ul and 0.5ug/ul respectively, into *Dsim w*^501^ and *IL43* which endogenously express Cas9 from the X chromosome. These strains were generated by introgressing the X chromosome from *y w* p{nos-Cas9, w^+^} in pBac{3XP3::EYFP, attp} sim 1087, which was kindly provided by David Stern (56). The cut site of two transgenic stocks were verified from each of the injected strains (Extended Data Fig. 5a and 5b). Transgenic *Dsim w*^501^ and *IL43* males heterozygous for the mutation were then crossed to non-injected *IL43* and *D. simulans w*^501^ virgins respectively. These crosses were amplified and the F1 males carrying the mutation (hemizygous for *trn* allele) were phenotyped as described previously (Extended Data Fig. 6c). For raw phenotypic data, see Supplementary File 2f.

In order to assess the effect of *trn* reciprocal hemizygotes to clasper phenotype, we first tested for normality and merged measurements from identical *trn* reciprocal hemizygotes (as described previously). We then conducted an ANOVA followed by a Tukey’s test for clasper bristle number, clasper area, tibia length and posterior lobe size between reciprocal hemizygotes (Supplementary File 2c).

## Supporting Information

Supporting information includes nine supplementary files.

## Author contributions

M.D.S.N., A.P.M., J.F.D.H. and P.G. designed the experiments. M.D.S.N., A.P.M. supervised and K.M.T., M.D.S.N., A.P.M. contributed reagents to the project. J.F.D.H., M.D.S.N., A.B. and C.C.M. performed the introgression mapping and RNAi experiments. J.F.D.H. and M.K carried out morphological analyses and SEM. K.M.T. performed the RNA-Seq experiments and analysis. A.P., J.F.J. and M.D.S.N. performed the *trn* sequence analysis. J.F.D.H. performed all other experiments. J.F.D.H. and M.D.S.N. analysed the data. M.D.S.N., A.P.M and J.F.D.H wrote the manuscript. All authors read and commented on the manuscript.

## Acknowledgements

We thank Christian Schlötterer, Christina Muirhead and Daven Presgraves for facilitating access to population genetic data. This work was funded by grants from the NERC (NE/M001040/1) and BBSRC (BB/M020967/1) to A.P.M., a JSPS KAKENHI (15J05233) grant to K.M.T. and a Genetics Society Summer Studentship grant to A.B..

---

**Extended Data Fig. 1.**
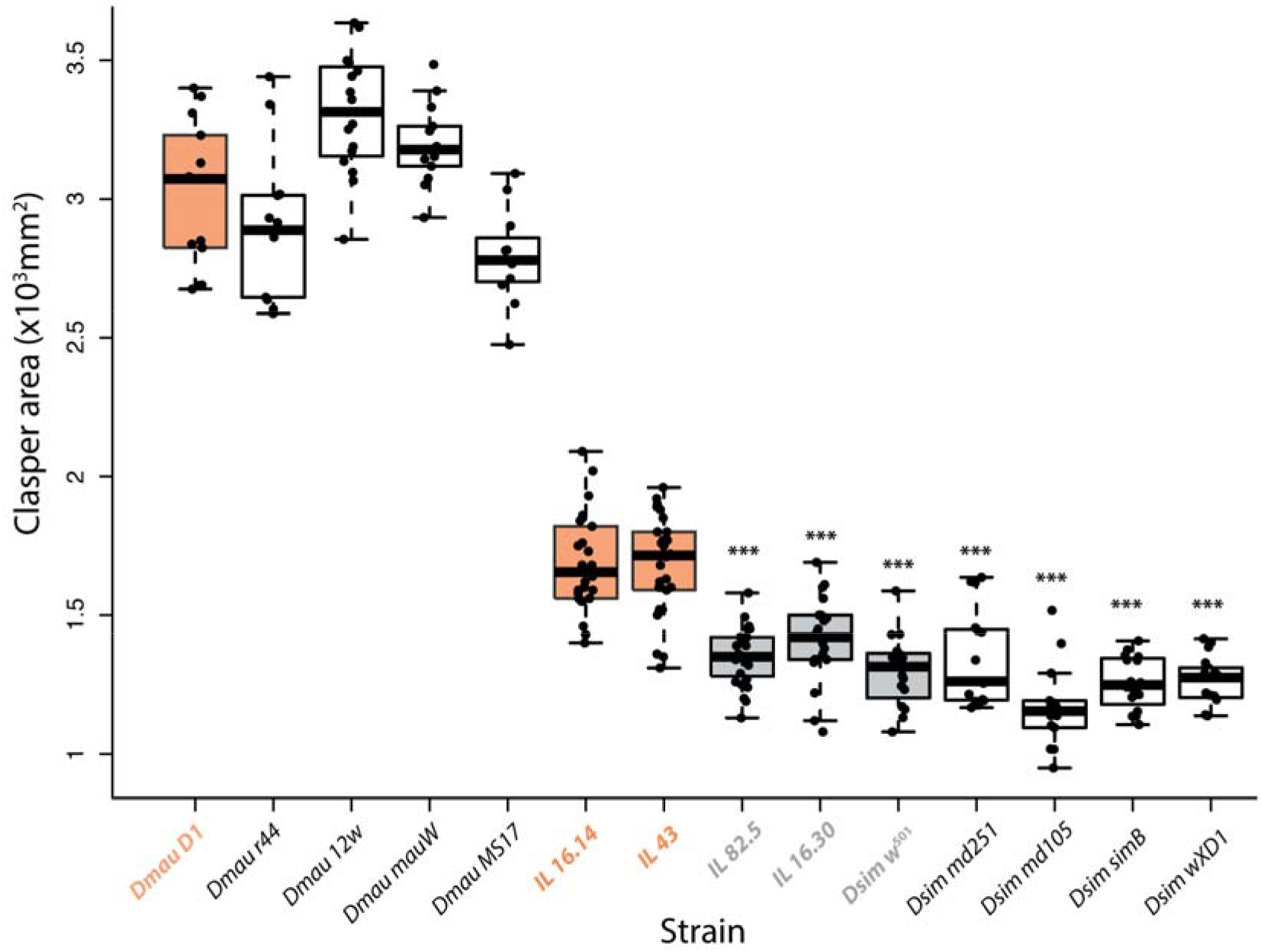
Intraspecific variation in clasper area and the effect of the C2 introgressed region. The parental strains used during introgression mapping, *Dmau D1* and *Dsim w*^501^, are shaded in orange and grey respectively, as well as the lines used to define the boundaries of C2 (with *IL 16.14* and *IL 43* in orange, containing *D. mauritiana* alleles over C2 and *IL 82* and *IL 16.30* in grey, containing *D. simulans* alleles). Both *Dsim w*^501^ and *Dmau D1* exhibit intermediate clasper size phenotypes that are within the species range. *** indicate *D. simulans* strains and ILs that differ significantly (*p* < 0.001) from *IL 16.14* and *IL 43*.

**Extended Data Fig. 2.**
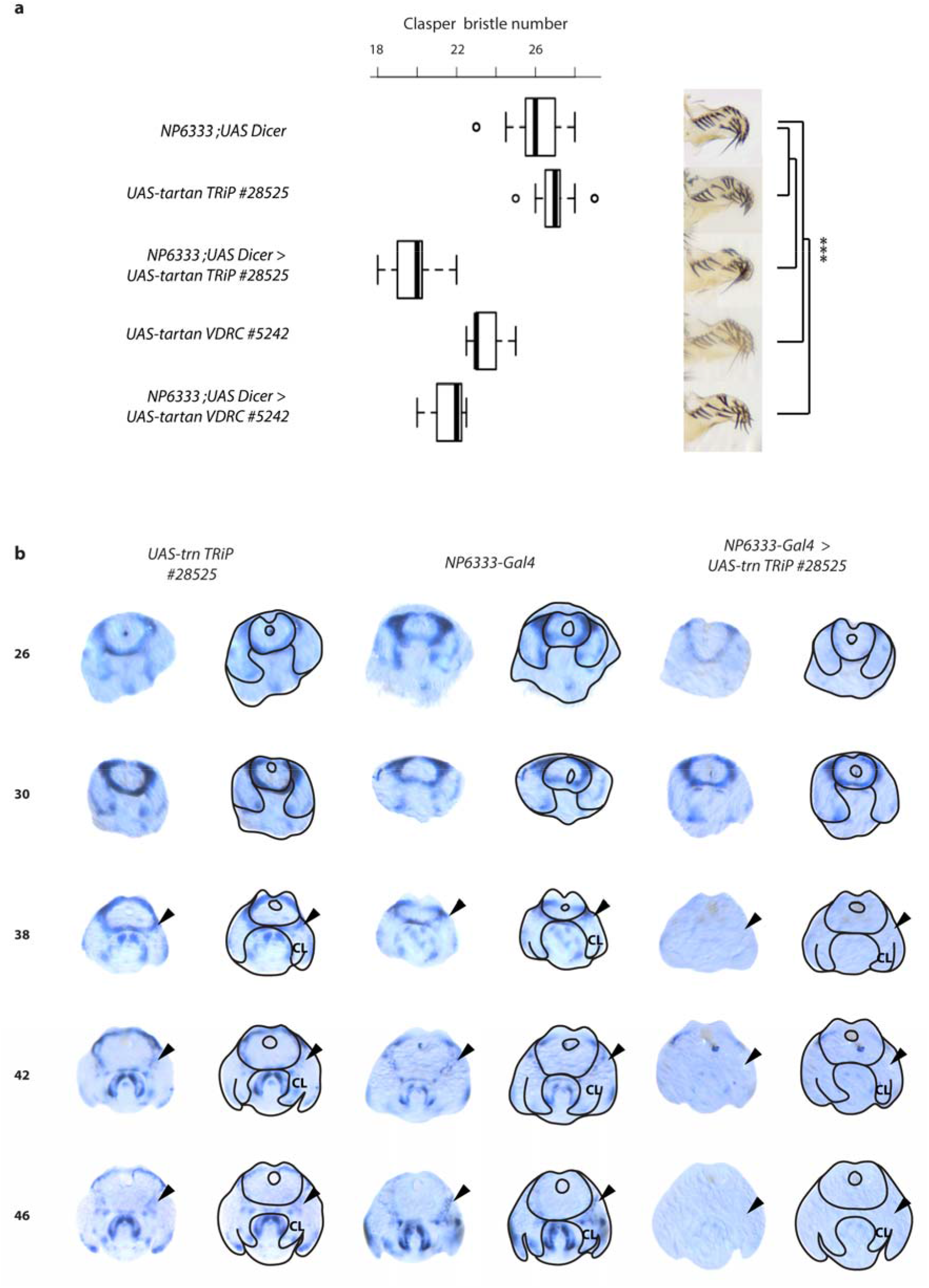
*tartan* RNAi in *D. melanogaster* male genitalia. **(a)** Effect on clasper bristle number of UAS-trn TRiP (#28525) or VDRC lines (#5242) combined with the NP6333 driver and UAS controls. Significant differences were detected between knockdowns and parental controls (#28525 *p* <0.001, F_(2, 52)_ = 211.1 and #5242 *p* < 0.001, F_(2, 39)_ = 153.9). Both constructs generated knockdown males that were significantly different from parental controls (\*\*\**p* < 0.001). **(b)** In situ hybridisation (using the #28525 UAS-trn TRiP line) on *trn* expression in developing claspers of *trn* knockdown compared to controls. Time in hAPF is indicated on the left. Black arrowheads indicate the base of the developing claspers. Boxes show the range, upper and lower quartiles, and the median for each sample.

**Extended Data Fig 3.**
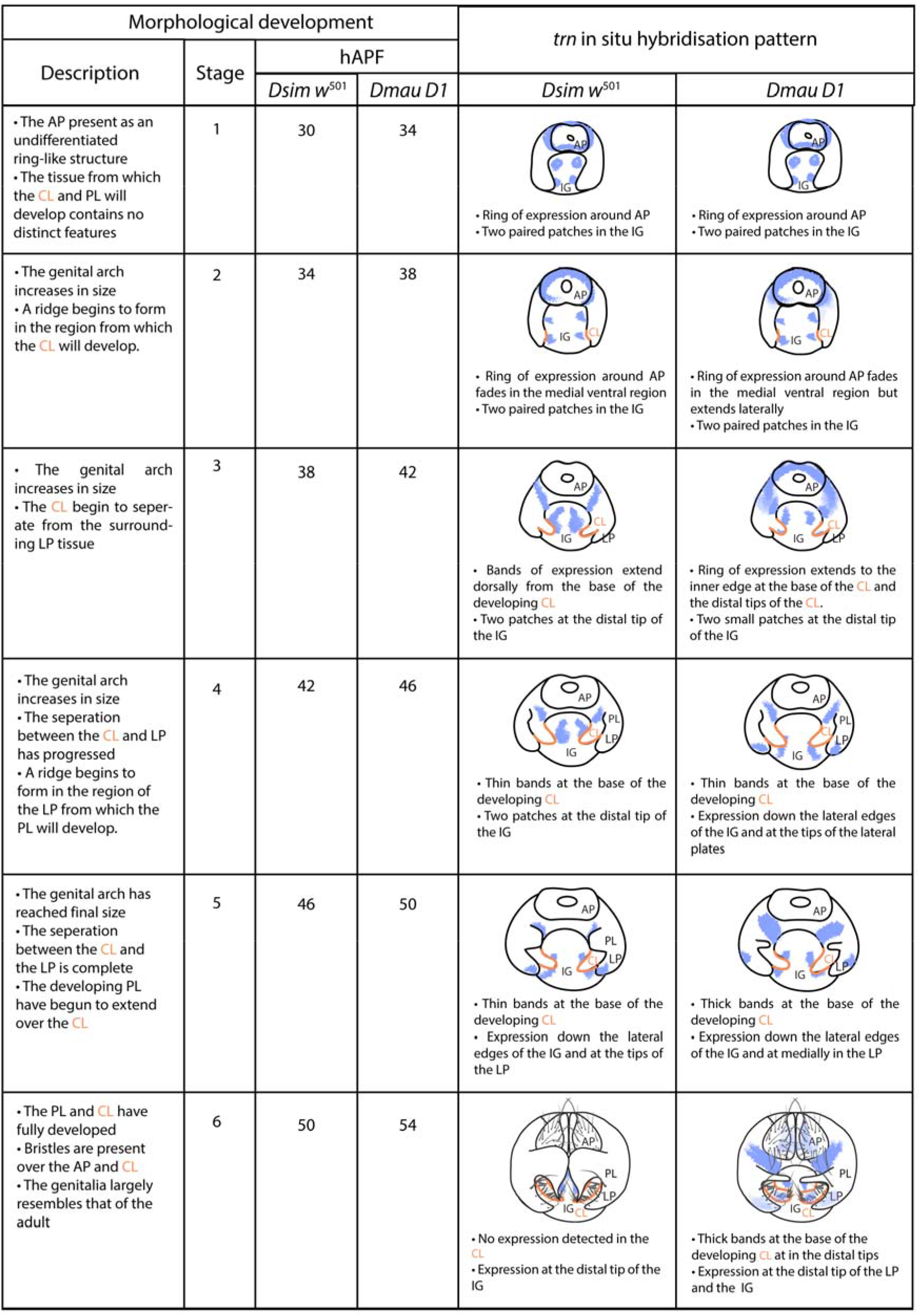
Temporal differences in male genital development and *trn* spatial expression between *D. simulans* and *D. mauritiana*. Based on morphological observations, *Dmau D1* male terminalia develops ~ 4 hours slower than *Dsim w*^501^. Many aspects of *trn* expression differ between the species; namely that *trn* is expressed in a broader domain, and for longer, at the base of the developing claspers in *Dmau D1* compared to *Dsim w*^501^.

**Extended Data Fig. 4.**
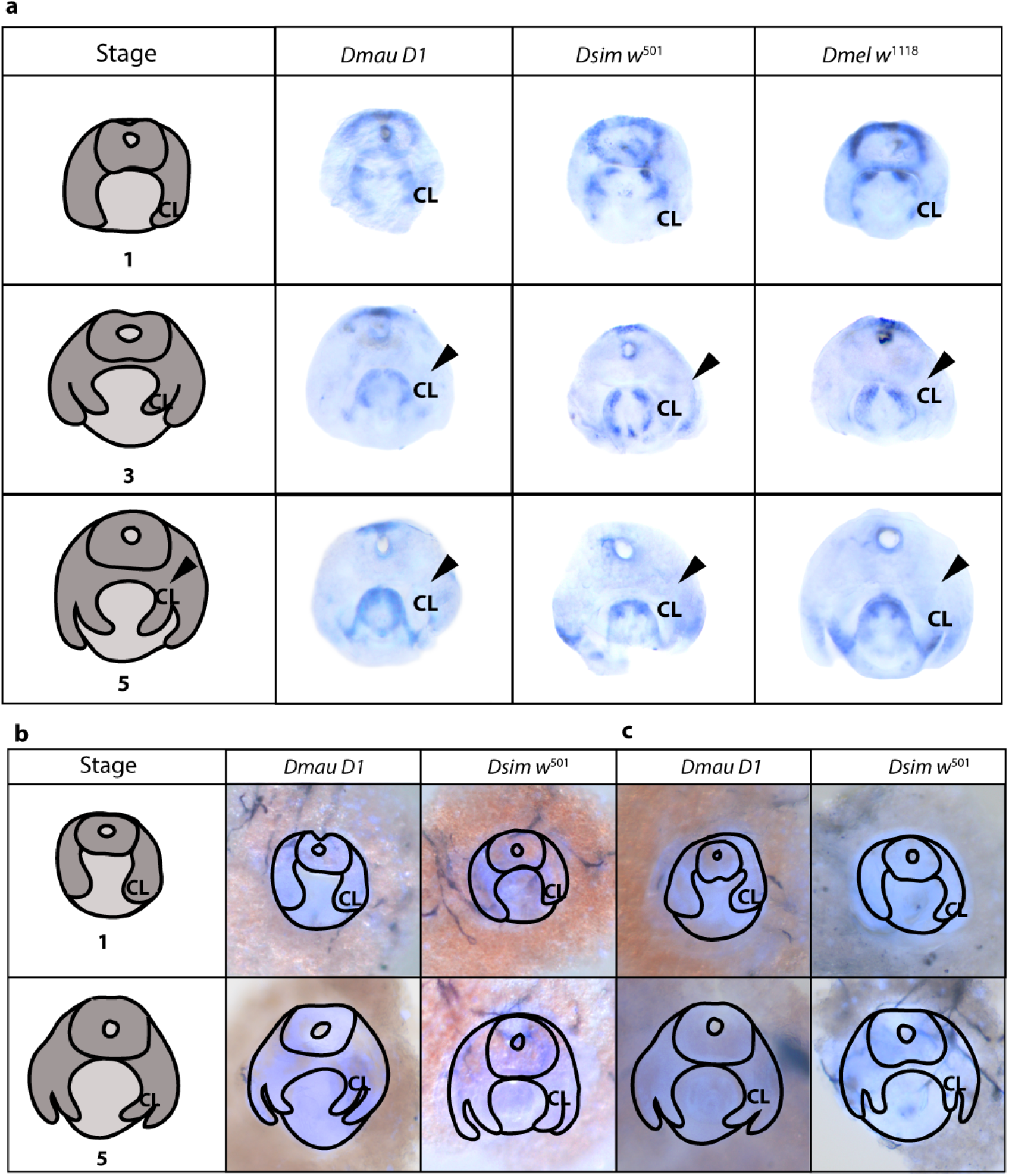
**(a)** Expression of *caps* in the developing male genitalia of *Dmau D1, Dsim w*^501^ and *Dmel* w^1118^, during early (Stage 1), middle (Stage 3) and late stages of development (Stage 5). Note that *caps* had no detectable expression at the base of the claspers (black arrowheads). We did not detect expression of *CG11279* **(b)** or *CG34429* **(c)** in claspers of *Dmau D1* and *Dsim w*^501^ during early (Stage 1), and late stages of development (Stage 5).

**Extended Data Fig. 5.**
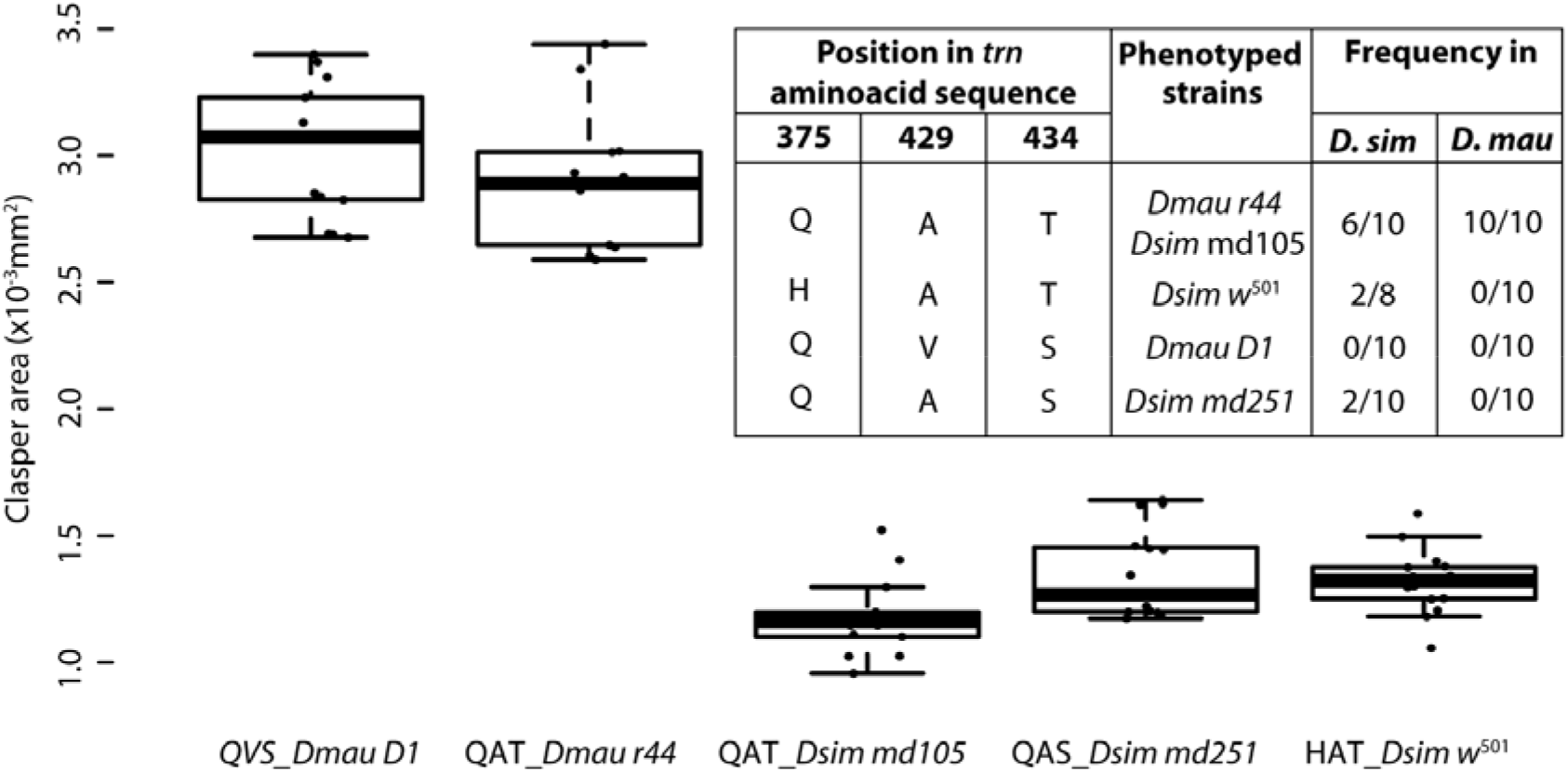
Clasper area of *D. simulans* and *D. mauritiana* strains carrying alternative amino acids at the three non-synonymous substitutions found between our mapped strains *Dsim w*^501^ and *Dmau D1*. Frequency information was obtained by resequencing the ten strains of each species kindly sent to us by the Presgraves lab. While we did not find strains carrying alternative versions of all possible combinations of the three non-synonymous substitutions, we were able to compare the clasper areas of strains that differed in each of these sites. The QAT combination, which only differs by one amino acid from that of *Dsim w*^501^, is the most common in both species, and is shared with *D. sechellia* and *D. melanogaster* (Supplementary File 6a). The clasper area of *Dmau r44*, which carries this amino acid combination, does not differ significantly from that of *Dmau D1* (*p* > 0.05, Supplementary File 2e) but differs from that of *Dsim md105*, which is also QAT (*p* < 0.001, Supplementary File 2e) by a similar degree to the difference between each of the *D. mauritiana* strains and *Dsim w*^501^ (Supplementary File 2e). This suggests that the H375Q substitution does not contribute to the clasper size difference between *Dmau D1* and *Dsim w*^501^. In addition, *Dsim md251*, which only differs in the second amino acid substitution from *Dmau D1* and only the third amino acid substitution from *Dmau r44*, differs from these *D. mauritiana* to the same extent as the other *D. simulans* strains, suggesting that A429V and T434S are also unlikely to contribute to the difference between the two mapped strains in this study (Supplementary File 2e). Clasper bristle number (not shown) yields similar results (Supplementary File 2e).

**Extended Data Fig. 6.**
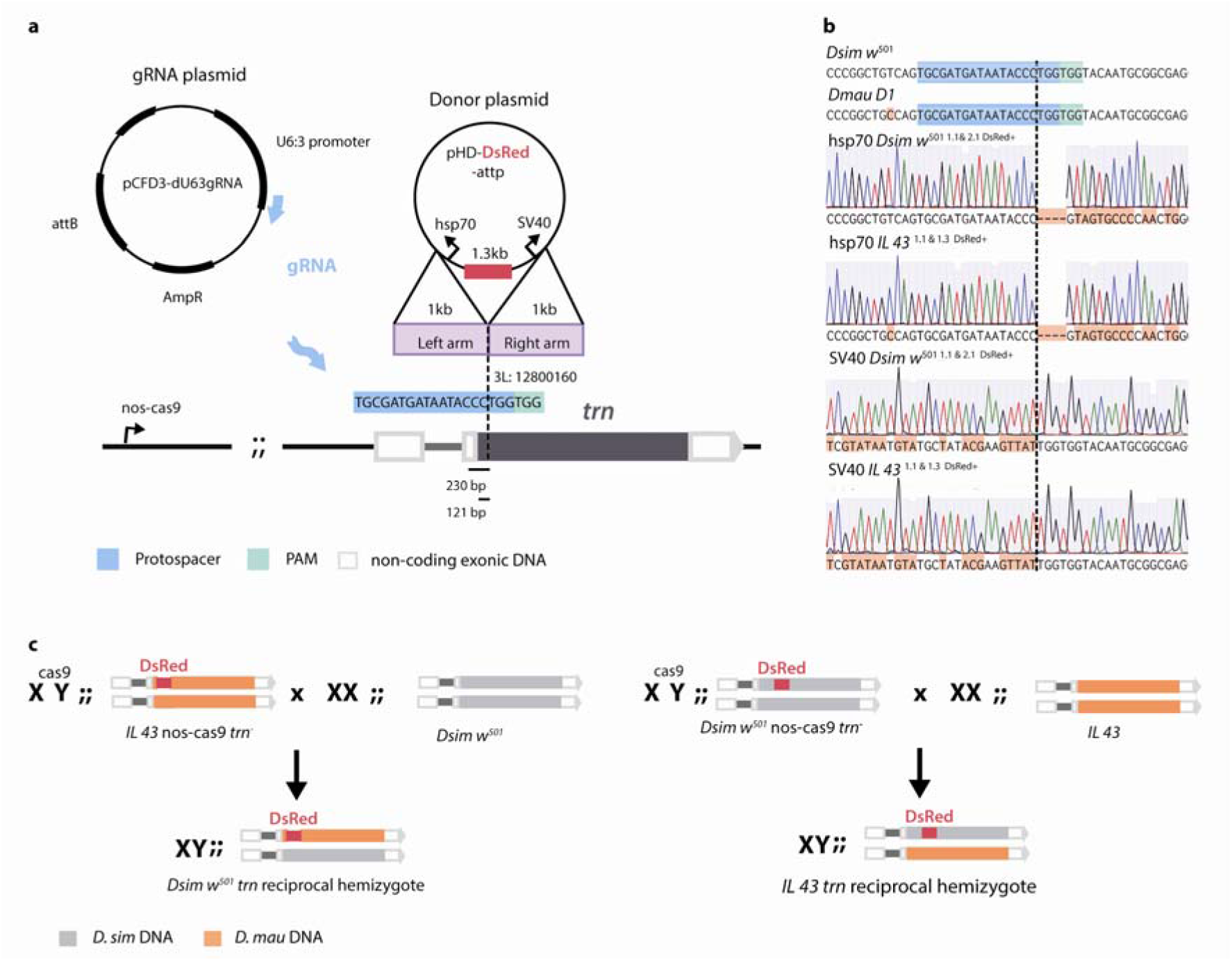
Strategy for generation of *trn* reciprocal hemizygotes of *Dsim w*^501^ and introgression line IL 43 (nanos-Cas9). **(a)** gRNA and donor plasmids used to insert DsRed into the coding region of trn using CRISPR-mediated homologous recombination. **(b)** Chromatograms illustrating the disruption of the *trn* reading frames in *Dsim w*^501^ and *IL 43*. **(c)** Crossing strategy to generate male reciprocal hemizygotes for *trn*.

